# Comparing bulk and single-cell methodologies and models to profile gene expression, chromatin accessibility and regulatory links in endothelial cells treated with TNFα

**DOI:** 10.64898/2026.03.13.711357

**Authors:** Jennifer Zevounou, Ken Sin Lo, Christopher S. McGinnis, Ansuman T. Satpathy, Guillaume Lettre

## Abstract

Genome-wide association studies (GWAS) have identified thousands of non-coding variants associated with complex traits and diseases. However, it remains challenging to pinpoint the causal genes that are regulated by associated genetic variants. Connecting causal non-coding variants with genes can rely on methods that identify direct physical interactions (e.g. chromosome conformation capture) or on probabilistic models that predict regulatory links. These statistical models take advantage of gene expression and chromatin accessibility profiles generated in cells and tissues by bulk or single-cell (sc) methodologies. Here, we tested whether using bulk or sc RNAseq/ATACseq data and corresponding predictive enhancer-to-gene models impact the prioritization of causal GWAS genes. Using non-treated and TNFα-treated human endothelial cells *in vitro* as a well-controlled experimental system, we show that bulk and sc RNAseq/ATACseq profiles are similar and highlight the same biology (e.g. biological pathways). Despite these similarities, we show using GWAS results for coronary artery disease (CAD) and diastolic blood pressure that applying enhancer-to-gene models designed for bulk or sc methodologies can yield differences in terms of captured heritability, fine-mapped variants and linked genes. For instance, at one CAD locus, the bulk-based ABC model predicts a regulatory link with *BCAR1*, whereas the sc-based model scE2G prioritizes a different gene (*CFDP1*). On the same experimental model, our results indicate that choosing between a bulk or sc approach will influence regulatory link model predictions; this should be considered when planning functional experiments to characterize GWAS discoveries.

## INTRODUCTION

Most genetic variants associated with complex human phenotypes by genome-wide association studies (GWAS) are non-coding and likely influence phenotypic variation by regulating gene expression (1). These genetic variants alter the activities of open chromatin regulatory sequences (e.g. enhancers), which in turn lead to dysregulation of gene expression. The identification of the genes controlled by distal regulatory elements in which GWAS variants reside is essential to gain molecular insights into human disease risk. Indeed, an understanding of the cell-type and developmental specificities of the links that connect regulatory elements and genes can lead to innovative genome editing strategies to treat human diseases (2,3). However, it is challenging to infer enhancer-to-gene interactions because physical distance alone is an imperfect predictor; some distal regulatory elements can act >100-kb from their targeted genes (4).

Chromosome conformation capture methods (e.g. 3C, 4C, HiC) have been developed to capture direct physical interactions between regulatory elements and genes (5). These methods are effective, but also technically challenging and costly, limiting their application in most laboratories. As an alternative solution, many statistical models have been developed to predict interactions between regulatory elements and genes (hereafter referred to as “regulatory links” which, unless specified, exclude promoters (6–9). These models take advantage of vast amount of gene expression, chromatin accessibility and chromosome 3D contact profiles in human cells and tissues to predict enhancer-to-gene interactions. For these models, the input data come mostly from bulk experiments, where the profiles are an average of multiple cells and cell-types. More recently, the characterization of gene expression and chromatin accessibility at single-cell (sc) resolution using multiome methods has enabled the development of newer probabilistic enhancer-to-gene models (10,11). While both bulk and sc enhancer-to-gene prediction models have been experimentally validated in a limited number of settings, it is currently unknown whether the predictions of the models are consistent when analyzing bulk and sc profiles from the same experimental system.

Vascular endothelial cells form the inner layer of blood vessels and have critical functions in controlling inflammatory responses, angiogenesis, the vascular tone and thrombosis (12). These cells are directly involved in the etiology of coronary artery disease (CAD) and hypertension, two of the most common cardiovascular diseases in the world. Partitioned heritability studies have revealed that open chromatin sites identified in vascular endothelial cells capture significant fraction of the GWAS signals for CAD and high blood pressure (13–15). Therefore, connecting non-coding GWAS variants with causal genes in vascular endothelial cells could improve the prevention, prediction and treatment of many cardiovascular diseases.

In this study, we analyze bulk and sc data from immortalized human vascular endothelial cells (teloHAEC) that are non-treated or stimulated with the pro-inflammatory cytokine TNFα. As expected, we find that RNAseq and ATACseq results from bulk and sc methods are largely consistent. However, when we apply bulk (ABC) and sc (scE2G) enhancer-to-genes models to teloHAEC gene expression and open accessibility profiles, we find that the models often link associated variants with different candidate genes. Our results can impact how to plan follow-up experiments to characterize GWAS loci in endothelial cells and other cell-types.

## RESULTS

### Measuring endothelial cell responses to TNF**α** stimulation

To compare the outputs of bulk and sc methods, we used as model immortalized human aortic endothelial cells (teloHAEC) non-treated (NT) or treated for 4 or 24 hours with the pro-inflammatory cytokine TNFα. TNFα treatment induces a robust and reproducible inflammatory response in teloHAEC that models the state of activated endothelial cells in the context of vascular diseases like hypertension or coronary artery disease (CAD). We previously used bulk RNAseq, ATACseq, histone H3 lysine 27-acetylated (H3K27ac) ChIPseq and HiC to characterize teloHAEC endothelial cell activation by TNFα^1^. This data is publicly available on NCBI GEO (GSE126200). For the sc component, we used new 10X multiome data generated in teloHAEC under the same TNFα conditions (**Methods**). The sc data is available from the IGVF Data Portal (https://data.igvf.org/analysis-sets/IGVFDS1583PWNS/). In this study, we compare gene expression, open chromatin peaks and links between genes and distal regulatory elements in teloHAEC that were measured or predicted with bulk and sc modalities.

### teloHAEC gene expression and open chromatin measurements are concordant between the bulk and single-cell methods

Gene expression levels measured by bulk and sc RNAseq were highly concordant (**Supplementary Tables 1-2**). Furthermore, in the three comparisons (TNFα 4hr vs NT, TNFα 24hr vs NT, TNFα 24hr vs TNFα 4hr), gene expression fold-changes were highly correlated between bulk and sc RNAseq (**Fig. 1A**, **Supplementary Fig. 1A-B**), despite many differentially expressed genes (DEG) being significant with only one RNAseq modality (**Fig. 1B**). Notably, more DEGs were identified using bulk RNAseq (N_bulk_=643 vs N_sc_=177), probably because the bulk RNAseq experiment was better powered to detect small fold-change differences (**Supplementary Fig. 1C-E**) or because of the differences between the statistical methods used to assess significance (**Methods**). Enrichment analyses showed that although DEG sets were only partially overlapping between bulk and sc RNAseq, the associated biological pathways were highly concordant (**Fig. 1C** and **Supplementary Fig. 2**).

**Figure 1.**
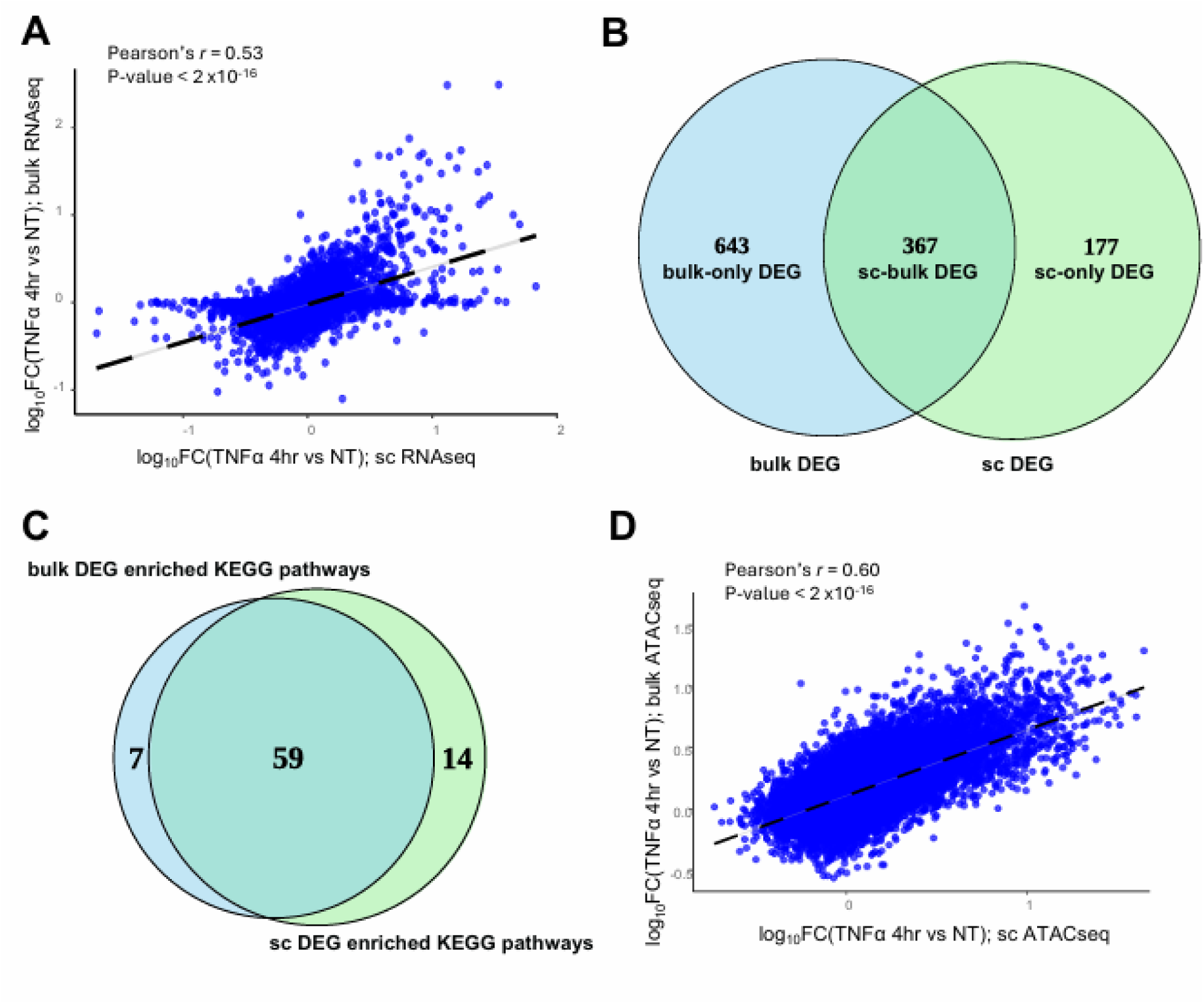
Bulk and single-cell RNAseq and ATACseq modalities generate concordant results. (**A**) Gene expression fold-changes (FC) measured using single-cell (sc) (x-axis) or bulk (y-axis) RNAseq are correlated when comparing treated (4 hours with the pro-inflammatory cytokine TNFα) and non-treated (NT) teloHAEC. (**B**) Among 11,221 genes measured by both sc and bulk RNAseq, we identified 1,187 differentially expressed genes (DEG), including 367 DEG identified by both methods. (**C**) Although many DEG are identified by only sc or bulk RNAseq, the list of enriched KEGG biological pathways is largely consistent when using DEG from sc or bulk RNAseq. (**D**) Open chromatin FC measured using sc (x-axis) or bulk (y-axis) ATACseq are correlated when comparing treated (4 hours with the pro-inflammatory cytokine TNFα) and NT teloHAEC.

To compare the bulk and sc ATACseq results (**Supplementary Table 3**), we first paired open chromatin sites between both modalities (**Supplementary Fig. 3** and **Methods**). Across the three conditions, we found 86,014 bulk-sc ATACseq peak pairs which involved 63% and 70% of the sc and bulk ATACseq peaks, respectively (**Supplementary Table 4**). Fold-change values were highly correlated between assays (**Fig. 1D** and **Supplementary Fig. 4A-B**). Similar to the RNAseq experiment, bulk ATACseq was able to detect differentially open peaks (DOP) with smaller fold-change differences (**Supplementary Fig. 4C-E**).

### Regulatory links between candidate cis-regulatory elements and genes in teloHAEC identified using bulk and single-cell data

Next, we tested whether the differences observed between the bulk and sc RNAseq/ATACseq results impacted predictions of regulatory links between gene transcriptions start sites (TSS) and candidate cis-regulatory elements (cCRE). We applied the activity-by-contact (ABC) model to predict regulatory links between cCRE and genes using genomic data derived from bulk experiments in teloHAEC NT or treated with TNFα (ATACseq, H3K27Ac ChIP-seq and HiC, **Methods**)(7). For comparison, we analyzed the teloHAEC multiome scRNAseq+scATACseq data and applied the scE2G model to infer regulatory cCRE-gene links in the sc dataset (11). The scE2G model is built on top of the ABC model and allows for the integration of sc-level information like correlation between cCRE accessibility and gene expression. The predicted bulk-(ABC) and sc-based (scE2G) cCRE-gene links are available in **Supplementary Tables 5-10.**

When we compared the bulk- and sc-inferred regulatory links, we noted that many of the best scoring cCRE-gene predictions based on the scE2G model were for promoters (**Fig. 2A-C**). In contrast, the ABC predictions were more distal to the gene bodies because promoters were excluded as potential distal regulatory elements (**Fig. 2A**). For each of the three conditions (NT, TNFα 4hrs and 24hrs), we paired regulatory links if the bulk/sc cCRE overlapped >250-bp and were connected to the same genes (**Supplementary Table 11**). In total across the three conditions, we found 35,464 pairs of regulatory cCRE-gene links predicted by the ABC and scE2G models, corresponding to 20-32% of all predicted links (**Supplementary Fig. 5**) and 15% of all regulatory regions (**Fig. 2D**). For downstream analyses, we compared the properties of the ABC-scE2G, ABC-only and scE2G-only regulatory regions that are predicted to be linked to genes in teloHAEC.

**Figure 2.**
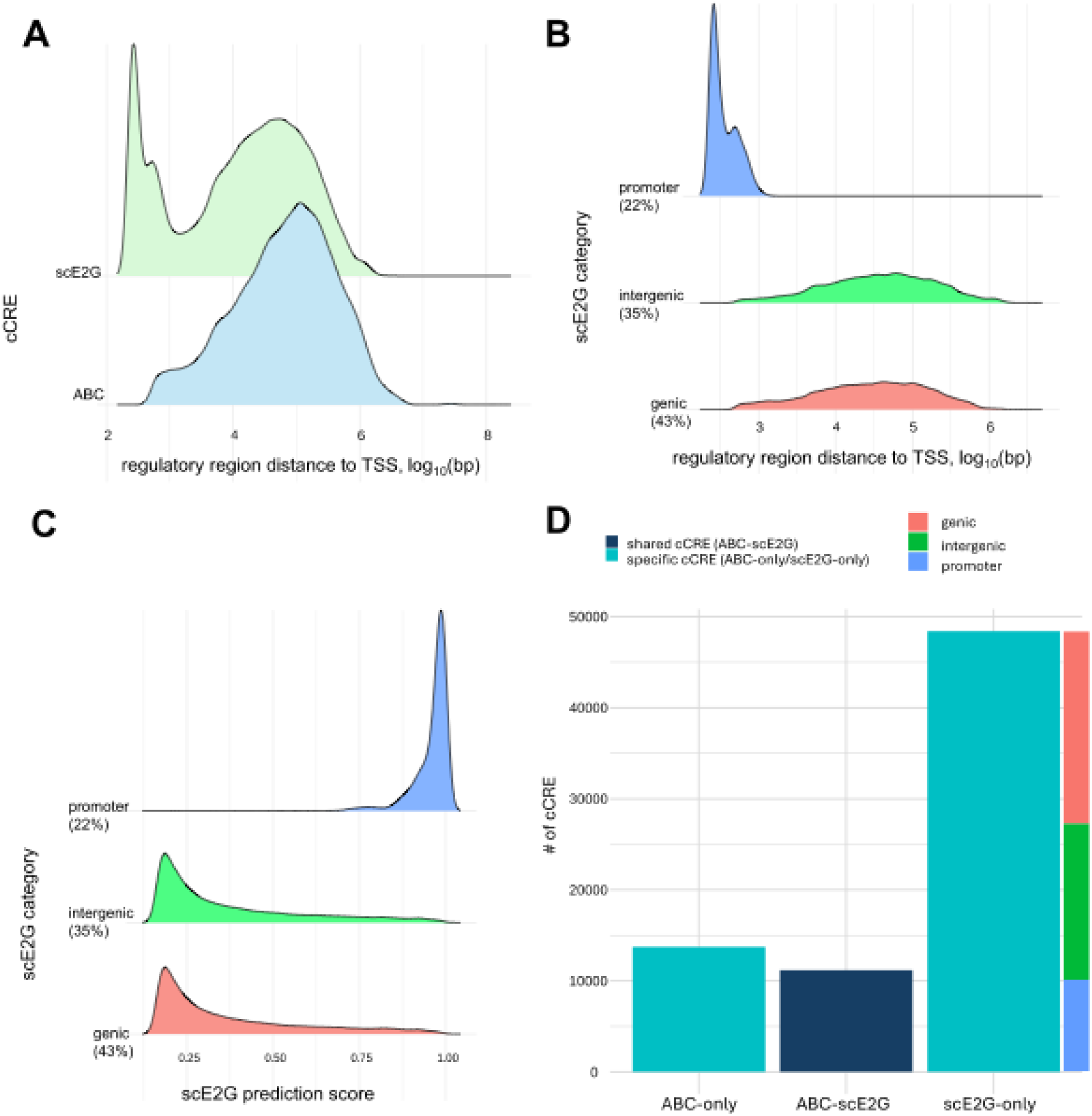
Identification of regulatory links between candidate cis-regulatory elements and genes in teloHAEC using sc and bulk data. (**A**) Distribution of the physical distance between candidate cis-regulatory elements (cCRE) and genes for links inferred using the ABC (bulk data) and scE2G (sc data) models. (**B**) Stratification of the scE2G predictions based on genomic annotations. As expected, the shorter regulatory links involve cCRE that are annotated as promoters. (**C**) High scE2G scores are mostly assigned when links are predicted between promoters and genes. (**D**) Bar plot summarising the number of cCRE identified with both the bulk and sc data (ABC+scE2G), bulk only (ABC-only) and sc-only (scE2G-only) across all treatment conditions. For sc-only cCRE, the number of promoters, intergenic and genic cCRE identified are also reported. TSS, transcription start site; bp, base pairs.

### teloHAEC cCRE-gene links are enriched for coronary artery disease and diastolic blood pressure heritability and fine-mapped variants

We and others have shown previously that the heritability for CAD and blood pressure (BP) is enriched among genetic variants located in open chromatin regions found in vascular endothelial cells (13–15). We took advantage of this aspect of the genetic architecture of these two phenotypes to compare the bulk- and sc-based cCRE-gene predictions. We used linkage disequilibrium (LD) score regression to partition the heritability of CAD and diastolic BP (DBP) among the endothelial cCREs associated with ABC-scE2G, ABC-only and scE2G-only link predictions (**Supplementary Tables 12-13**)(16). We found that mostly all cCRE subsets captured a significant fraction of the heritability for these two cardiovascular phenotypes (**Fig. 3A-B**). Unexpectedly, the ABC-only cCRE showed a non-significant enrichment for CAD which, upon further analyses, appear to be due to instability in the LD score regression jackknife estimate (16). Indeed, excluding a single block resulted in a significant enrichment estimate for CAD in ABC-only cCRE (**Supplementary Fig. 6A-B)**.

**Figure 3.**
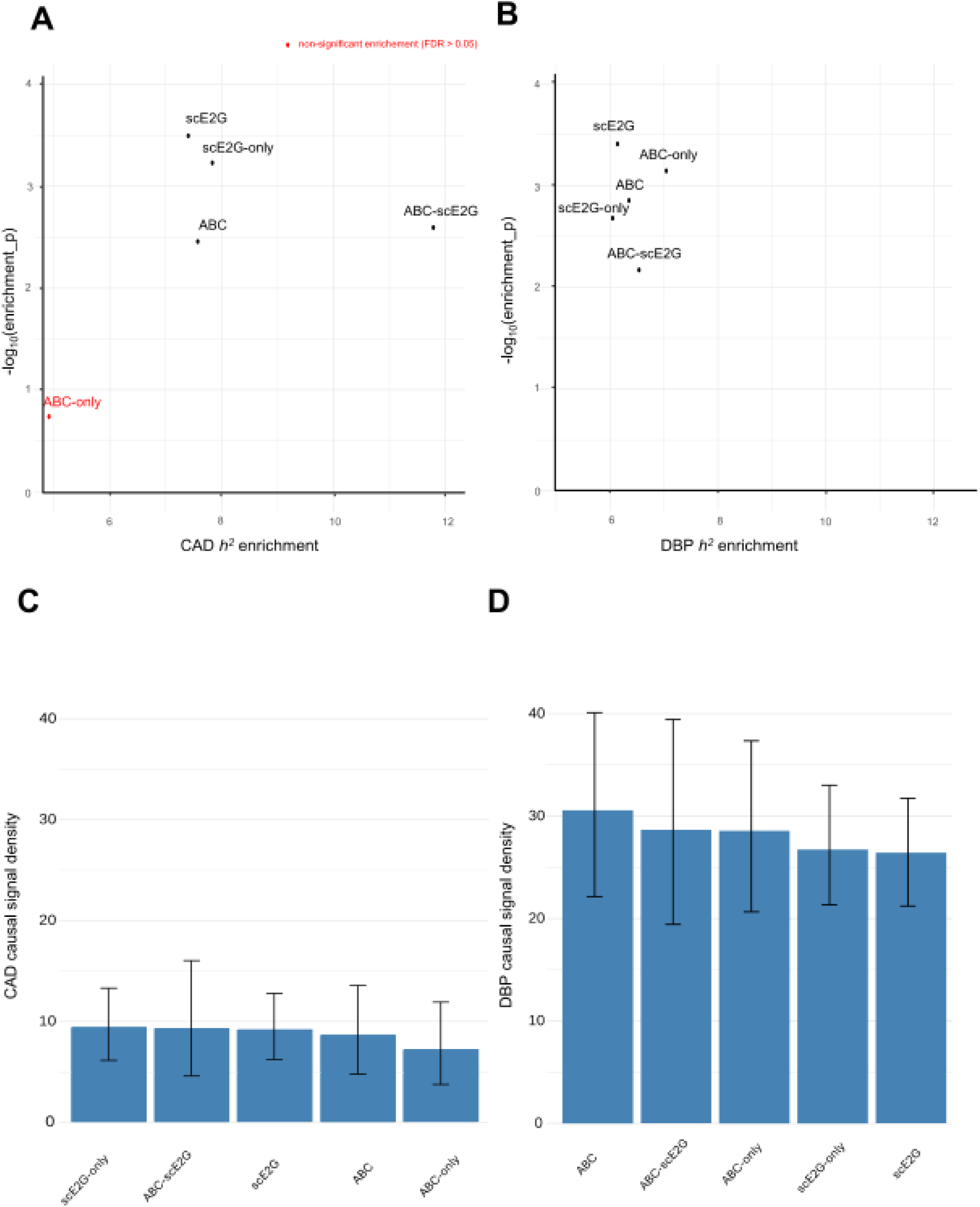
Coronary artery disease (CAD) and diastolic blood pressure (DBP) heritability estimates, and fine-mapped variants captured by teloHAEC regulatory links. Scatter plots of (**A**) CAD and (**B**) DBP linkage disequilibrium (LD) score regression-based heritability estimates (x-axis) for variants within five regulatory link categories, along with their corresponding enrichment P-values (y-axis). All enrichment estimates are significant (false discovery rare [FDR] <0.05) except for ABC-only (red). (**C-D**) Bar plot of CAD and DBP causal signal density calculated for five different categories of regulatory links. We define the causal signal density as the sum of the posterior inclusion probabilities for all fine-mapped variants that overlap with the regulatory elements in each category (**Methods**). We calculated the 95% credible intervals by bootstrapping.

To determine how well cCRE implicated in regulatory links capture genome-wide significant association results, we fine-mapped the CAD and DBP GWAS summary statistics using RSparsePro (**Supplementary Tables 14-15**)(17). Then, we calculated the sum of the posterior inclusion probability (sumPIP) for all variants that belong to 95% credible sets and that overlap with cCRE (**Methods**). Finally, because the number of predictions and size of the regulatory regions vary between bulk and sc (**Supplementary Table 16**), we corrected sumPIP by the proportion of the genome covered by the cCRE subset to obtain a “causal signal density” (**Fig. 3C-D** and **Supplementary Fig. 6C-D**). Using the causal signal density results, we concluded that all cCRE groupings implicated in predicted regulatory links capture a similar amount of the CAD (**Fig. 3C**) and DBP (**Fig. 3D**) causal association signals, and the regulation of gene expression in endothelial cells is particularly suitable to study the genetic architecture of DBP (**Fig. 3C-D**, average causal signal densities for CAD and DBP are 8.7 and 28.2, respectively).

### Prioritization of CAD candidate causal genes using teloHAEC regulatory links and Open Targets predictions

Open Targets has compiled multiple lines of evidence, including GWAS results, to prioritize genes implicated in complex human diseases and traits (18). We used the Open Targets “association scores” for CAD to assess whether the teloHAEC ABC or scE2G cCRE-gene links connected more regulatory regions with candidate causal CAD genes. For this analysis, we focused on links between an Open Targets CAD gene and a cCRE that includes CAD fine-mapped variants (**Supplementary Table 17**). A similar number of CAD genes were linked by ABC-only, scE2G-only and ABC-scE2G predictions despite differences in the number of regulatory cCRE-gene links (**Fig. 4A**, dark green). The enrichment of connected CAD:non-CAD genes was higher for predicted links found by both ABC and scE2G (10-times) and lower for ABC-only links (**Fig. 4A**). The Open Targets CAD association scores were higher for the candidate genes that were connected to cCRE, independently of the methods used to predict regulatory links (**Fig. 4B**). When we excluded promoters from scE2G predictions, we found that the Open Targets association scores were marginally higher for candidate CAD genes connected by ABC-scE2G links (**Fig. 4B**). We found 59 (e.g. *IL6R, FES*) and 44 (e.g. *BTBD16, DHX36*) candidate CAD genes predicted by Open Targets that were only linked to fine-mapped GWAS variants by the scE2G and ABC models, respectively (**Fig. 4C** and **Supplementary Tables 18-19**). In contrast, we only found 66 CAD candidate genes that were linked by both the ABC and scE2G models (e.g. *TGFB1, NOS3, PLPP3*)(**Fig. 4C** and **Supplementary Tables 18-19**).

**Figure 4.**
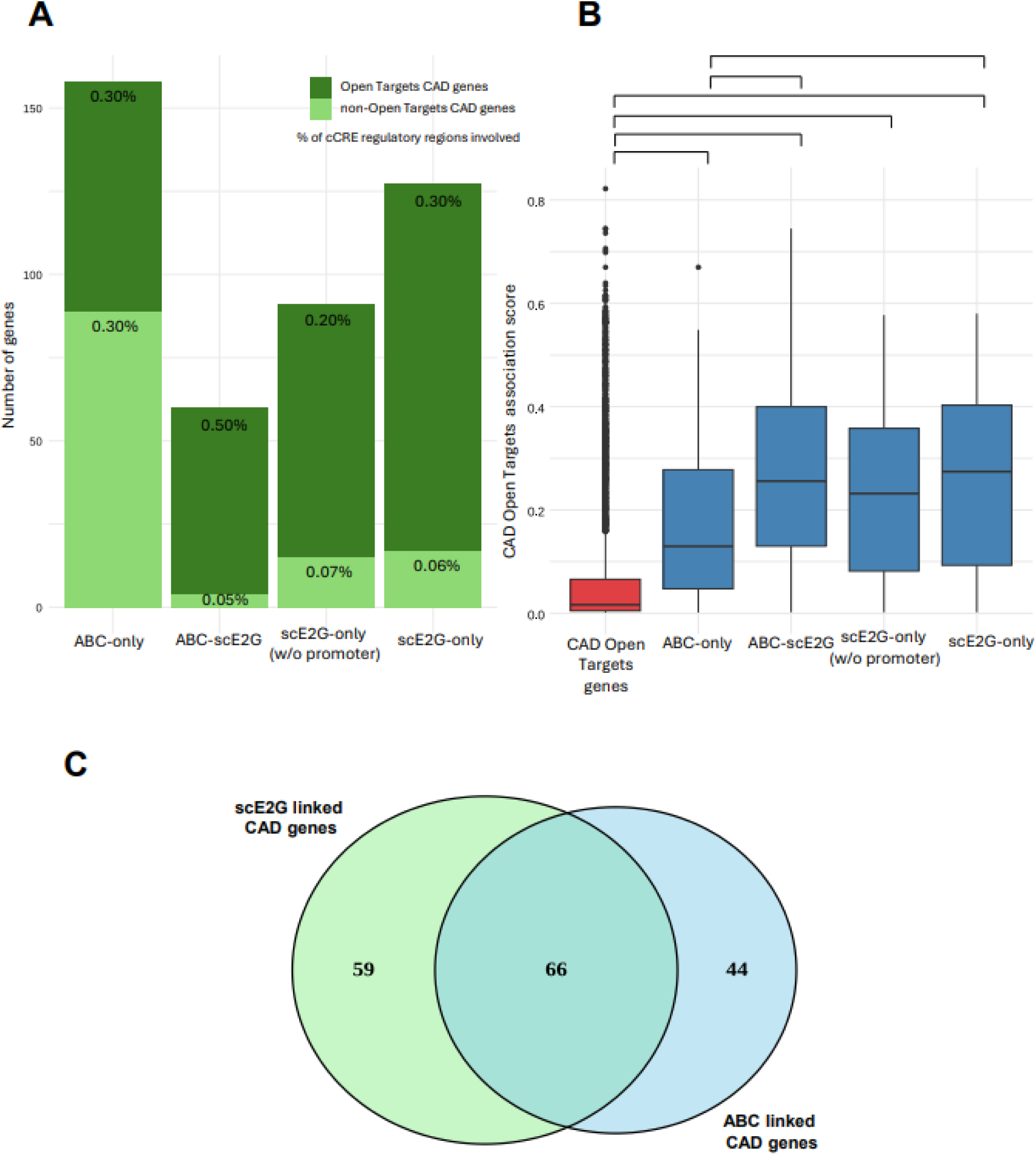
Connecting coronary artery disease (CAD)-associated variants and Open Targets CAD candidate causal genes in teloHAEC. (**A**) Bar plot of the number of genes linked to candidate cis-regulatory elements (cCRE) in five different categories of regulatory links. The dark green fraction corresponds to the proportion of CAD causal genes predicted by Open Targets. The percentage of cCRE linking causal and non-causal CAD genes are reported. (**B**) Distribution of Open Targets CAD candidate causal gene association scores for genes linked to regulatory elements through different categories. The box in red shows the distribution of Open Targets association scores for all CAD genes, independently of connections to regulatory elements. In blue are the distributions of the Opent Targets association scores for CAD genes linked to cCRE through different enhancer-to-gene models. The scores for the distributions in blue are significantly higher than the scores for any of the distributions in red (Wilcoxon’s *P*-value<0.05). The distribution of the Open Targets scores for ABC-only are significantly lower than the scores for ABC-scE2G (Wilcoxon’s *P*-value=6.6×10^-3^) and scE2G-only (Wilcoxon’s *P*-value=7.3×10^-3^). (**C**) Venn diagram of the number of CAD causal genes predicted by Open Targets and linked by either or both the scE2G and ABC models. The list of genes in each category is in **Supplementary Table 19**.

To illustrate examples of gene regulation that may impact CAD risk, we selected three loci with candidate CAD genes expressed in endothelial cells, and visualized open chromatin sites and predicted regulatory links. At the *TGFB1* locus, results from bulk and sc methodologies were consistent, prioritizing the same GWAS variants linked to the *TGFB1* TSS (**Fig. 5A**). TGF-β signaling can promote the angiogenic potential of endothelial cells (19). *IL6R* encodes the receptor of the pro-inflammatory cytokine interleukin-6 (IL6), which is known to accelerate the process of atherosclerosis (20). Although the chromatin accessibility profiles at the *IL6R* locus were similar when comparing bulk and sc ATACseq results, only the scE2G model predicted links that implicated fine-mapped CAD variants in the regulation of *IL6R* expression (**Fig. 5B**). Finally, at the *BCAR1*/*CFDP1* locus, we found three fine-mapped variants (chr16:75377623 A>T, chr16:75377721 A>G, chr16:75378364 C>T) in a cCRE identified by bulk and sc ATACseq that was linked to the *CFDP1* TSS by scE2G but with the *BCAR1* TSS by ABC (**Fig. 5C**). Altogether, these results and examples suggest important differences between bulk and sc strategies to connect non-coding regulatory variants from GWAS with candidate causal genes.

**Figure 5.**
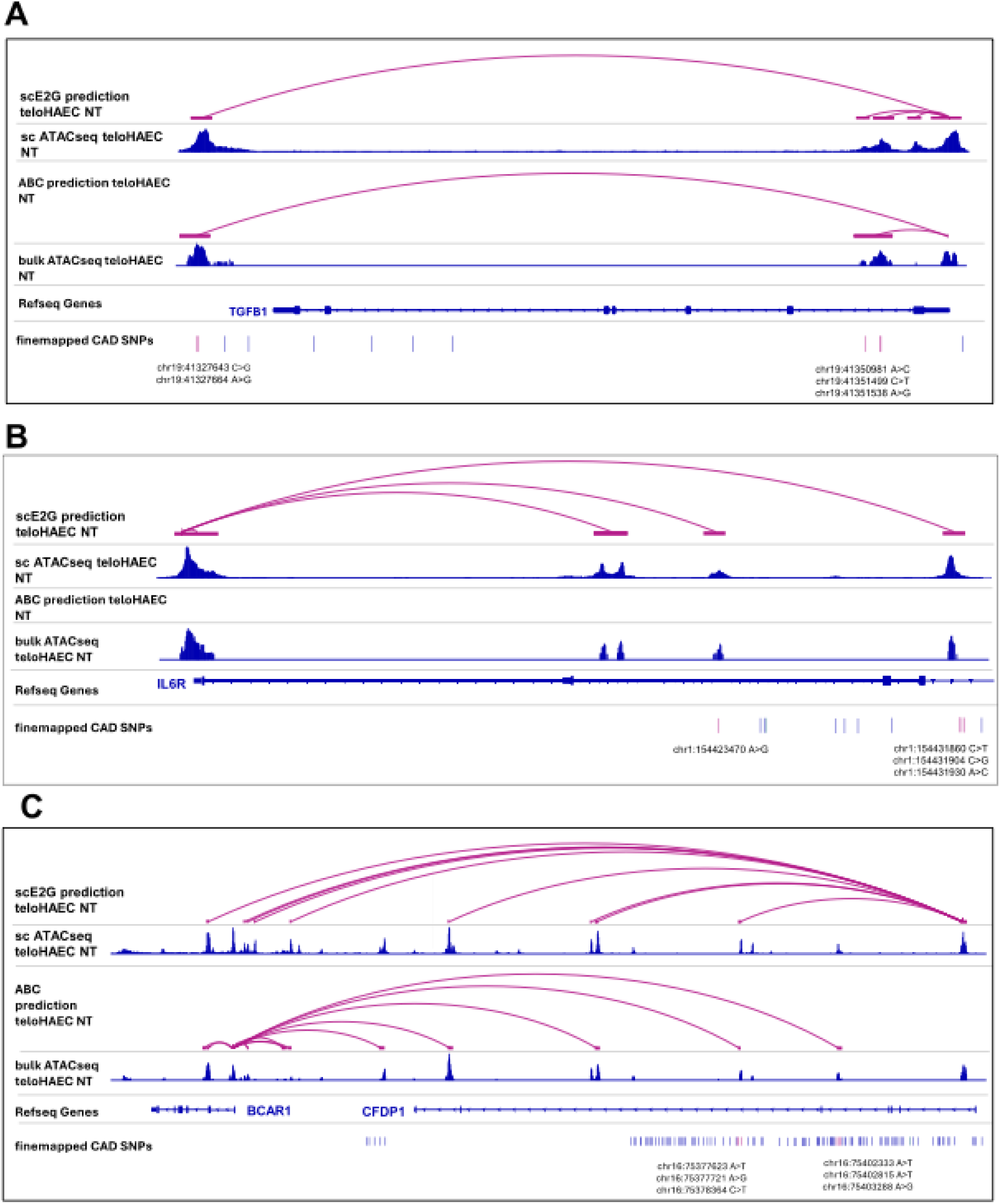
Examples of regulatory links identified by scE2G and/or ABC at different coronary artery disease (CAD) loci. Genomic representations of the (**A**) *TGFB1*, (**B**) *ILR6* and (**C**) *BCAR/CFDP1* CAD-associated loci. In each panel, we included the regulatory link predictions, the ATACseq peaks (all genes are expressed in teloHAEC) and the fine-mapped CAD variants. Variants overlapping cCRE are highlighted in pink. These panels represent three different scenarios: (**A**) both ABC and scE2G models identify the same cis-regulatory element (cCRE) for *TGFB1*, (**B**) only one model (scE2G) identifies a cCRE connected to *IL6R* and (**C**) both models identify cCRE but link them to different target genes: *CFDP1* for scE2G and *BCAR1* for ABC.

## DISCUSSION

The identification of the causal genes that are regulated by non-coding genetic variants associated with human diseases and traits is an important step in translating human genetic findings to clinical insights. This task is difficult because regulatory elements do not necessarily control the expression of the closest genes. Several statistical models have been developed to predict regulatory links between open chromatin elements and genes using as input data generated with bulk or sc methodologies. Sc-multiome data offers the double advantage to profile single cell-types from mixtures of cells found in tissues, and to measure gene expression and chromatin accessibility in the same cells at the same time. However, when compared with bulk methods, sc methods have less sensitivity to capture genes expressed at low levels or chromatin peaks that are less accessible (**Supplementary Fig. 1 and 4**). In our study, we show that these methodological differences can impact the predictions of state-of-the-art enhancer-to-gene predictive models.

Vascular endothelial cells are highly relevant to the etiology of cardiovascular diseases, and GWAS have implicated endothelial functions in CAD, stroke and hypertension risk (13,21–23). Using gene expression and chromatin accessibility data from a well-characterized human vascular endothelial cell system, we compared the impact of the methods (bulk vs sc) and statistical models on enhancer-to-gene predictions. Despite working with a simple in vitro model of vascular endothelial cells, we found that predictions by the ABC and scE2G models were not always consistent across the same GWAS loci. Indeed, while regulatory links identified by both the ABC and scE2G models connected fine-mapped variants with excellent candidate genes at many loci (e.g. *TGFB1, NOS3, PLPP3*), we also found many candidate genes linked to variants by only the ABC (e.g. *BTBD16, DHX36*) or the scE2G model (e.g. *IL6R, FES*)(**Supplementary Tables 18-19**). We even identified multiple examples where the same fine-mapped variant was linked to different causal genes by the ABC and scE2G models (**Supplementary Table 19**). These examples include cases where the linked genes are next to each other (e.g. *BCAR1* and *CDFP1*, *COL4A1/2* and *RAB20*, **Fig. 5C**, **Supplementary Table 19**), far from each other (e.g. chr7:140049546 A>G is linked to *SLC37A3* by ABC [284-kb] but *TBXAS1* by scE2G [29-kb], where only the latter is a CAD gene), or with different CAD Open Targets association scores (e.g. chr1:113907040 C>T is linked to *AP4B1* by ABC [Open Targets score=0.1] but *DCLRE1B* by scE2G [Open Targets score=0.47]). These results indicate that the source of the data and the models used to connect non-coding GWAS variants and genes can strongly influence how to plan functional experiments.

Our study has several limitations. First, we only tested two of the many models that exist to connect regulatory elements and genes. However, we selected the ABC and scE2G models because they outperform most of the other models in specificity/sensitivity analyses (7,11). Second, we compared model predictions from a single experimental system. As paired bulk-sc data become available, it will be interesting to determine if our conclusions from the analysis of vascular endothelial cells also apply to other cell-types. Finally, our results are dependent on the quality and completeness of available cell profiles. As methods become more comprehensive to measure expression and chromatin accessibility (in particular sc methods), predictive models will undoubtedly improve.

While 3D contacts are more easily measured in a bulk setting, using prediction models based on sc multiome measurements allow to estimate the variability in gene expression and chromatin accessibility between cells. Integrating both approaches may therefore provide a more robust framework for predicting cCRE-gene links (24). In summary, we show that the profiling methods used to make enhancer-to-gene predictions can influence the list of predicted causal genes from GWAS. Our results emphasize the importance to consider (or combine) alternative gene prioritization strategies (e.g. HiC, expression/protein quantitative trait loci [eQTL/pQTL]) before planning functional experiments to characterize GWAS discoveries.

## METHODS

### Bulk dataset

Bulk RNAseq, ATACseq and HiC data from immortalized human endothelial cells (teloHAEC; non-treated, treated for 4 and 24 hours with TNFα) have been described before (13) and are available from NCBI GEO: www.ncbi.nlm.nih.gov/geo/query/acc.cgi?acc=GSE126200.

### Single-cell multiome datasets

Sequencing libraries were constructed using the 10X Genomics Single Cell ATAC and RNA Multiome kit and the MULTI-seq lipid oligo hashing protocol for sample labelling before pooling. Sequencing was performed using Illumina NovaSeqX. FASTQ files were processed using Cell Ranger (cell-ranger (6.0.0)(25), cellranger-atac (2.0.0)(26)) and deMULTIplex2 (1.0.2)(27). Joint gene expression and chromatin accessibility profiles were obtained for 18,977 cells, totaling approximately 83 million RNA-seq UMI and 248 million ATAC-seq fragments. Thesc-multiome data is available on the IGVF Data Portal (https://data.igvf.org/analysis-sets/IGVFDS1583PWNS/). Quality-control (QC), normalization and visualization of the sc-multiome RNA and ATAC teloHAEC dataset was performed using Seurat (5.3.0)(28) and Signac (1.15)(10) in R (4.4.0). We first selected high quality cells using the following thresholds: 1600< nCount_RNA <16000, mitochondrial DNA <30%, number of genes >1100, log_10_GenesPerUMI >0.87, 3000< nCount_ATAC <40000, FriP >0.45, blacklist_ratio <1e-03, nucleosome signal <0.65, TSS enrichment >3.5. Genes present in less than 10 cells were removed from the analysis. Doublets were identified and removed using scDblFinder (1.20.2)(29). Gene expression normalisation and dimension reduction was performed using the functions SCTransform() and RunPCA() in Seurat, respectively. For scATACseq, RunTFIDF() and RunSVD() were used for peak accessibility normalisation and dimension reduction. Finally, scRNA and scATAC modalities were integrated together using the FindMultiModalNeighbors() function.

### Differential gene expression analysis

We used the differential gene expression results from Lalonde *al*. for the bulk teloHAEC data (13). For the scRNAseq dataset, we performed differential expression analysis using the Wilcoxon Rank Sum test implemented in FindMarkers(), comparing expression across treatments (NT, TNFα 4hr, TNFα 24h). Differentially expressed genes (DEG) were identified in the scRNAseq data using the following criteria: an absolute log_10_(fold-change [FC]) >0.3, a Bonferroni corrected P-value <0.05, and the gene tested being expressed in at least one percent of the cells (pct.1 >= 0.01 or pct.2 >=0.01). For the comparison of the bulk and sc DEG results, we focused on gene that were present in both analyses. Consistency in the direction and magnitude of gene expression change was assessed by performing Pearson’s correlation tests between sc and bulk estimated log_10_(FC).

### Pathway enrichment analysis

Pathway analysis was performed online on the sc and bulk datasets using DAVID (https://davidbioinformatics.nih.gov/)(30). Pathway enrichment was performed on all DEG identified, no matter the treatment condition, using all genes that passed QC steps, as background. Pathways with a nominal enrichment P-value <0.05 are described as significantly enriched.

### Differential chromatin accessibility analysis

Differential chromatin accessibility in the sc dataset was performed using the same tools and same criteria as in the differential gene expression analysis. Chromatin accessibility analysis was performed independently for the sc and bulk datasets. Peaks analysed in the sc and bulk datasets have different coordinates and therefore needed to be paired up before comparing chromatin accessibility results. We first compared the size of the scATACseq and bulk ATACseq peaks before selecting a 250-bp minimum overlap threshold to identify peaks present in both the sc and bulk datasets; we referred to these overlaps as sc-bulk ATACseq peak pairs. Results from the sc and bulk differential chromatin accessibility were compared for these peak pairs only. Comparison was done using the same methodology as for differential gene expression analysis comparison.

### scE2G regulatory links predictions

Regulatory links were inferred from the sc-multiome dataset using the scE2G model (v1.0), a genome wide predictor of enhancer-gene link which only requires multimodal scRNAseq and scATACseq data as inputs (11). We derived regulatory links for each treatment conditions (NT, TNFα 4hr, TNFα 24hr). As required by the model, we provided raw scRNAseq count matrix as well as scATAC fragment files for each treatment clusters. Following recommendation from Sheth et *al*. we filtered out regulatory links with a score <0.164(11). We ran scE2G using model multiome_powerlaw_v2.

### ABC regulatory links predictions on bulk data

We used the ABC model to predict regulatory enhancer-to-gene links between regulatory elements and genes (7). We reformatted the HiC contact data using Juicebox tools (31). The processed HiC matrices are then fit to a power-law decay model, which estimates how contact frequency decreases with genomic distance. These parameters allow the ABC pipeline to scale HiC interaction scores appropriately across the genome. In the first ABC step, we identified candidate enhancer regions by extending bulk ATACseq peak summits and filtering them against genomic blacklists and gene-proximal whitelist regions. In the second step, the pipeline integrates multiple datasets—ATACseq, H3K27ac ChIP-seq, gene annotations, and a curated list of ubiquitously expressed genes—to define regulatory “neighborhoods” around each gene. These neighborhoods group candidate enhancers with their nearby genes based on genomic proximity and chromatin activity. The third step uses both the enhancer lists and gene lists along with HiC contact maps to compute ABC scores for all candidate enhancer–gene pairs. The model then predicts which enhancers are likely to regulate each gene by applying a probability threshold (0.2).

### Comparions of regulatory links identified between genes and elements

We explored whether the regulatory links predicted by the scE2G model based on sc-multiome data were similar to those predicted by the ABC model, which relies on bulk chromatin accessibility measurement (H3k27ac ChIP-seq) and contact matrix (HiC). Three set of regulatory links, one per condition (NT, TNFα 4h, TNFα 24hr) were available for each method. Regulatory regions identified in the sc and bulk datasets have different coordinates and therefore needed to be paired up for comparison. Regulatory regions with an overlap >250-bp and linked to the same gene, identified with bedtools (2.31.0) pairToPair function, were paired up(32). Our downstream analysis of the scE2G and ABC predictions focused more on the regulatory regions than the links. For this reason, we simplified the regulatory region count by merging their coordinates across all treatment condition using bedtools merge function.

### Heritability enrichment analysis

We compared the CAD and DBP partitioned heritability enrichment in different subsets of the scE2G (sc) and ABC (bulk) predicted regulatory regions using S-LDSC method (v1.0)(16). Variants from the European 1000 Genome Phase 3 Project were annotated for their presence in each regulatory region subset, and their LD scores were re-calculated. Heritability enrichment in each regulatory region set was estimated individually using the baselineLD model (1000G_Phase3_baselineLD_v2.2).

### Fine-mapping of genome-wide association study (GWAS) results

We used RSparsePro (https://github.com/zhwm/RSparsePro_LD) to fine-map the CAD and DBP summary statistics (https://www.ebi.ac.uk/gwas/studies/GCST90310295, https://www.ebi.ac.uk/gwas/studies/GCST90132314)(33,34). First, the “get lead” script was run to identify loci of lead variants to fine-map, after removing variants with minor allele frequency (MAF) <0.00001, P-value >5e-8, or located inside the HLA locus (hg19:chr6:27477797-34448354). Variants that were seen in less than 90% of the total number of individuals were also filtered out. To calculate LD matrices, we used a subset of 20,000 unrelated and self-declared White British individuals from the UK Biobank. For each lead variant, an LD matrix was computed using PLINK (--matrix --r) for all variants with a minor allele frequency (MAF) >0.00001 located <500-kb from the lead variant. The “format ss” script from the RSparsePro package was run to ensure that alleles are properly aligned between the variants in the LD matrix and the variants in the GWAS summary statistics. Finally, the “rsparsepro ld” script was run to find credible sets that include at least 95% of total posterior inclusion probability (PIP). To compare the enrichment of causal GWAS variants located in predicted regulatory elements, we calculate the “causal signal density” (CSD), which we defined as the sum of the PIP for all fine-mapped variants located within a regulatory element set divided by the genomic size (i.e. number of base pairs) in that set.

### Identification of CAD candidate causal genes

Genes associated to CAD in the Open Target platform (18) were labelled as CAD candidate causal genes. For each regulatory region sets, we filtered out regions which did not map a CAD fine-mapped variant. We then listed the genes linked to the remaining regions and counted how many of them were CAD candidate causal genes. In Open Target, each gene is given an association score reflecting the number and strength of the evidence linking this gene to the phenotype of interest. We therefore compared the distribution of Open Target CAD candidate causal gene association scores between each regulatory region sets.

### Data visualisation

R plots were generated using base R functions, ggplot2(3.5.2)(35) and Venn Diagram(1.7.3)(36), ComplexUpset(1.3.3)(37). Regulatory regions gene-links were visualized with IGV (2.16)(38).

## DATA AVAILABILITY

The bulk data discussed in this publication have been deposited in NCBI’s Gene Expression Omnibus and are accessible through GEO Series accession number GSE126200 (https://www.ncbi.nlm.nih.gov/geo/query/acc.cgi?acc=GSE126200). The sc-multiome data is available on the IGVF Data Portal (https://data.igvf.org/analysis-sets/IGVFDS1583PWNS/). For fine-mapping, we used imputed genetic data from the UK Biobank (Project #62518).

## CODE AVAILABILITY

Scripts to analyze the data and draw figures are available at: https://github.com/jenniferzev/BULK_SC_PROFILE_TELOHAEC_TNFA

## ACKNOWLEDGEMENTS

This work was funded by the Montreal Heart Institute Foundation, the Joseph C. Edwards Foundation, the Canada Research Chair Program, the Canadian Institutes of Health Research (Project #168902), and the NIH/NHGRI Impact of Genomic Variation on Function Consortium (UM1HG012010) to G.L.. J.Z. acknowledges support from the Faculty of Medicine of the Université of Montréal (Bourse de mérite doctorale). C.S.M. is a METAVivor Early Career Investigator and acknowledges support from the NIH (1K99CA293137-01A1). A.T.S. acknowledges support from the NHGRI Impact of Genomic Variation on Function Consortium (UM1HG012076) as well as the Burroughs Wellcome Fund, Parker Institute for Cancer Immunotherapy, Pew Charitable Trusts, Cancer Research Institute, American Society of Hematology, and Baxter Foundation. Sequencing was performed at the UCSF Center for Advanced Technology, and we appreciate their team for technical support and access to computational resources that aided in data processing and alignment.

## AUTHOR CONTRIBUTIONS

J.Z. and G.L. designed the experiments and planned the analyses. J.Z. performed all analyses and generated the figures. K.S.L. performed the fine-mapping of the GWAS results. C.M. generated the sc-multiome data. G.L. and A.T.S. secured funding and supervised the work. J.Z. and G.L. wrote the manuscript with contributions from all authors.

## COMPETING INTERESTS

C.S.M. holds patents related to MULTI-seq. A.T.S. is a founder of Immunai, Cartography Biosciences, Santa Ana Bio, and Arpelos Biosciences, an advisor to 10x Genomics and Wing Venture Capital, and receives research funding from Astellas and Northpond Ventures. All other authors declare that they have no competing interests.

## Notes

https://github.com/jenniferzev/BULK_SC_PROFILE_TELOHAEC_TNFA

### Summary of Updates

The funding section of the manuscript was updated (J.Z.).

