## Supplementary Tables and Figures for "Comparing bulk and single-cell methodologies and models to profile gene expression, chromatin accessibility and regulatory links in endothelial cells treated with TNFα"

**Supplementary Table 1.** Results of the differential expression analysis in the scRNAseq dataset. See Excel file.

**Supplementary Table 2. Gene expression correlation tests.** Spearman’s correlation test between gene expression measured by bulk and sc RNAseq. For bulk and sc RNAseq gene expression values, we used mean Fragments Per Kilobase of transcript per Million mapped reads (FPKM) and mean unique molecular identifier (UMI), respectively.

|  | **Non-treated** | **TNFα 4hr** | **TNF α 24hr** |
| --- | --- | --- | --- |
| **Spearman’s**  **coefficient** | 0.8320748 | 0.8261277 | 0.829889 |
| **P-value** | <2.20e-16 | <2.20e-16 | <2.20e-16 |

**Supplementary Table 3.** Results of the differential chromatin accessibility analysis in the scRNAseq dataset. See Excel file.

**Supplementary Table 4. ATACseq peak pairs and counts.** Number of individual ATACseq peaks in each dataset and number of peaks involved in a sc-bulk ATACseq peak pair. The length of bulk ATACseq peaks being on average larger than those of scATAC peaks, one bulk ATACseq peak can be involved in multiple sc-bulk ATACseq peak pairs.

| **Description** | **Count** |
| --- | --- |
| Total number of scATAC peaks | 135,894 |
| Total number of bulk ATAC peaks | 95,490 |
| Number of sc-bulk ATAC peak-pairs | 86,014 |
| Number of scATAC peaks involved in at least one sc-bulk ATAC peak pair | 86,013 |
| Number of bulk ATAC peaks involved in at least one sc-bulk ATAC peak pair | 67,271 |

**Supplementary Table 5.** Predicted regulatory links using non-treated teloHAEC single-cell data (scE2G model). Coordinates are on build GRCh38. See Excel file.

**Supplementary Table 6.** Predicted regulatory links using teloHAEC (4hr, TNFalpha) single-cell data (scE2G model). Coordinates are on build GRCh38. See Excel file.

**Supplementary Table 7.** Predicted regulatory links using teloHAEC (24hr, TNFalpha) single-cell data (scE2G model). Coordinates are on build GRCh38. See Excel file.

**Supplementary Table 8.** Predicted regulatory links using non-treated teloHAEC bulk data (ABC model). Coordinates are on build GRCh38. See Excel file.

**Supplementary Table 9.** Predicted regulatory links using teloHAEC (4hrs, TNFalpha) bulk data (ABC model). Coordinates are on build GRCh38. See Excel file.

**Supplementary Table 10.** Predicted regulatory links using teloHAEC (24hrs, TNFalpha) bulk data (ABC model). Coordinates are on build GRCh38. See Excel file.

**Supplementary Table 11.** Regulatory links identified in teloHAEC that are predicted by the bulk (ABC) and single-cell (scE2G) models. The treatment (non-treated, 4hrs TNFalpha, 24hrs TNFalpha) is available in the last column. See Excel file.

**Supplementary Table 12.** Partitioned heritability results for CAD. Prop. SNPs: proportion of SNP inside de regions, Prop. h2(_std_error): proportion of heritability explained (standard error), Enrichment: Prop. h2/Prop. SNPs, Enrichment_std_error: enrichment standard error, Enrichment: Enrichment p-value.

| cCRE category | Prop._SNPs | Prop._h2 | Prop._  h2_std_error | Enrichment | Enrichment_std_error | Enrichment_p |
| --- | --- | --- | --- | --- | --- | --- |
| ABC | 0.00698438 | 0.052898711 | 0.015418882 | 7.573859232 | 2.207623598 | 0.003491908 |
| scE2G | 0.01566239 | 0.115939137 | 0.027032811 | 7.402390885 | 1.725969704 | 0.000318949 |
| ABC-only | 0.005527281 | 0.027155159 | 0.016834432 | 4.912932706 | 3.045698724 | 0.183659285 |
| scE2G-only | 0.014070418 | 0.110213422 | 0.027025709 | 7.832988374 | 1.920746684 | 0.000588897 |
| ABC-scE2G | 0.004538547 | 0.053490714 | 0.015316302 | 11.78586779 | 3.374714855 | 0.00254628 |

**Supplementary Table 13.** Partitioned heritability results for DBP. Prop. SNPs: proportion of SNP inside de regions, Prop. h2(_std_error): proportion of heritability explained (standard error), Enrichment: Prop. h2/Prop. SNPs, Enrichment_std_error: enrichment standard error, Enrichment: Enrichment p-value.

| cCRE category | Prop._SNPs | Prop._h2 | Prop._  h2_std_error | Enrichment | Enrichment  std_error | Enrichment_p |
| --- | --- | --- | --- | --- | --- | --- |
| ABC | 0.00698438 | 0.044366566 | 0.011691004 | 6.352255356 | 1.673878525 | 0.001400944 |
| scE2G | 0.01566239 | 0.096154376 | 0.022143598 | 6.139189008 | 1.413807071 | 0.0003905 |
| ABC-only | 0.005527281 | 0.038933143 | 0.009890514 | 7.043814869 | 1.789399599 | 0.000717402 |
| scE2G-only | 0.014070418 | 0.08511193 | 0.022651292 | 6.048997926 | 1.60985207 | 0.002092773 |
| ABC-scE2G | 0.004538547 | 0.029674916 | 0.009174226 | 6.538417738 | 2.021401692 | 0.006750085 |

**Supplementary Table 14.** RSparsePro results for CAD. See Excel file.

**Supplementary Table 15.** RSparsePro results for DBP See Excel file.

**Supplementary Table 16.** Number and size of the candidate cis-regulatory elements (cCRE) implicated in regulatory links using different methods or their intersection.

| **Methods to define regulatory links** | **Number of regions** | **Coverage (in kb)** |
| --- | --- | --- |
| ABC | 18,006 | 21,911 |
| scE2G | 53,118 | 48,622 |
| ABC-only | 13,729 | 17,291 |
| scE2G-only | 48,408 | 43,556 |
| ABC-scE2G | 11,174 | 14,442 |

**Supplementary Table 17.** Number of cis-regulatory elements (cCRE) implicated in regulatory links and that overlap with fine-mapped coronary artery disease (CAD)-associated variants.

| **Methods to define regulatory links** | **Number of regions** | **Number of SNP overlapped** |
| --- | --- | --- |
| ABC-only | 55 | 98 |
| scE2G-only | 156 | 226 |
| scE2G-only (w/o promoters) | 107 | 150 |
| ABC-scE2G | 57 | 96 |

**Supplementary Table 18.** CAD genes that are linked to cCRE in teloHAEC. See Excel file.

**Supplementary Table 19.** List of CAD fine-mapped variants linked to different genes by the ABC and scE2G model. Coordinates are on build GRCh38. See Excel file.

**SUPPLEMENTARY FIGURES**

**Supplementary Figure 1. Bulk and single-cell RNAseq results in teloHAEC cells are concordant.** Gene expression fold-changes (FC) are correlated when comparing single-cell (sc) (x-axis) and bulk (y-axis) RNAseq results. (**A**) teloHAEC TNFα 24 hours vs non-treated (NT) teloHAEC and (**B**) teloHAEC TNFα 24 hours vs teloHAEC TNFα 4 hours. Given the specificities of our RNAseq experiments, the bulk RNAseq protocol detects differentially expressed genes (DEG) with smaller FC in teloHAEC: (**C**) TNFα 4 hours vs NT, (**D**) TNFα 24 hours vs NT and (**E**) TNFα 24 hours vs TNFα 4 hours. abs, absolute value.

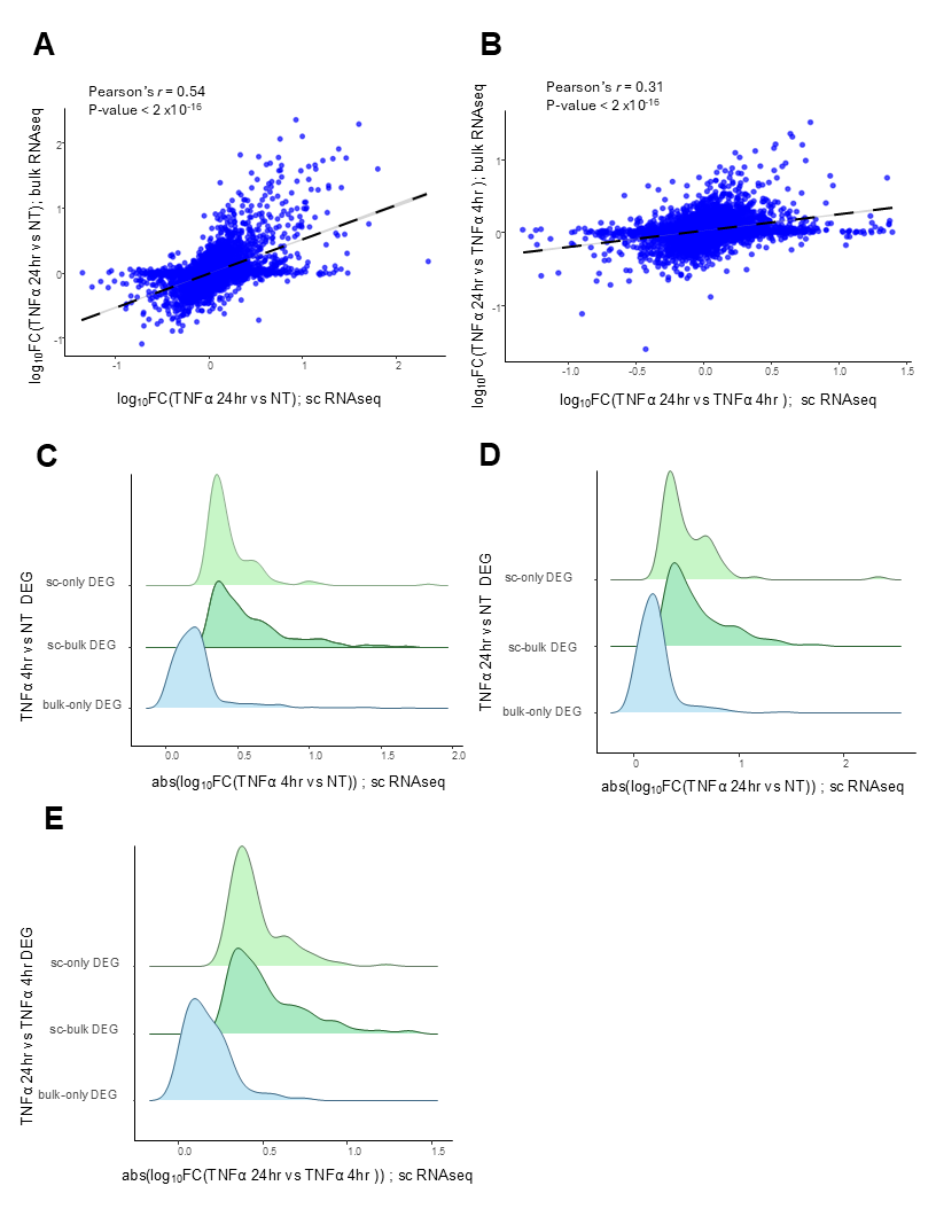

**Supplementary Figure 2.** **Bulk and single-cell RNAseq identify most of the same biological pathways in teloHAEC after TNFα stimulation.** Biological pathways that are enriched for: (**A**) both sc and bulk RNAseq differentially expressed genes (DEG), (**B**) only scRNAseq DEG and (**C**) only bulk RNAseq DEG. We note that most of the relevant biological pathways to endothelial functions and coronary artery disease (e.g. “fluid shear stress and atherosclerosis”, “lipid and atherosclerosis”, “TNF signalling pathway”) are identified by both scRNAseq DEG and bulk RNAseq DEG (nominal *P*-value<0.05).

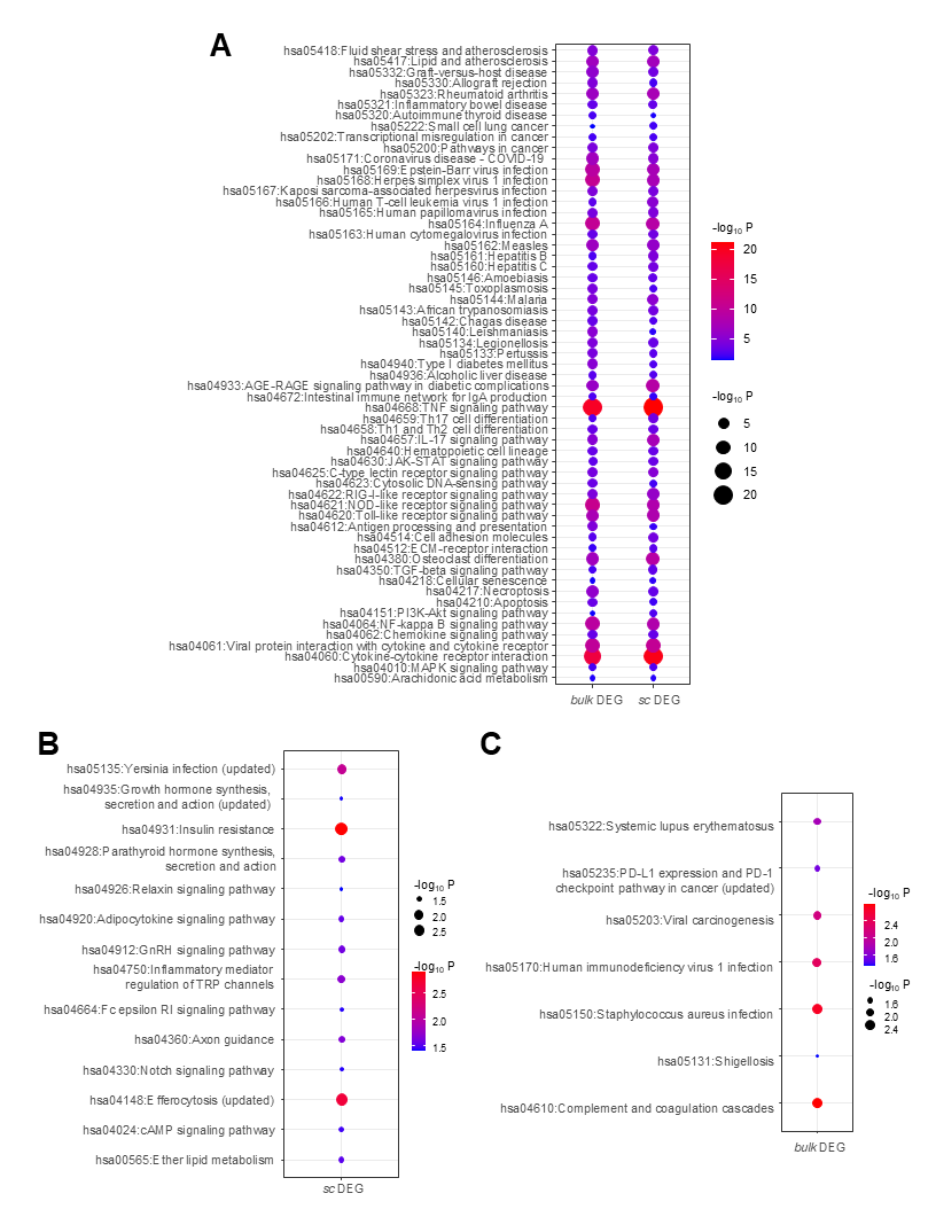

**Supplementary Figure 3. ATACseq-defined open chromatin peak lengths in teloHAEC.** (**A**) The open chromatin peaks identified by bulk ATACseq are on average larger than the scATACseq peaks, which are 450-500-bp long. (**B**) To pair open chromatin peaks identified by bulk and sc ATACseq, we required a 250-bp overlap. Most scATACseq peaks (~450-500-bp) are completely embedded within the larger bulk ATACseq peaks. For **A** and **B**, ATACseq peaks in teloHAEC from all treatment conditions are included (non-treated, TNFα 4hr, TNFα 24hr). bp, base pairs.

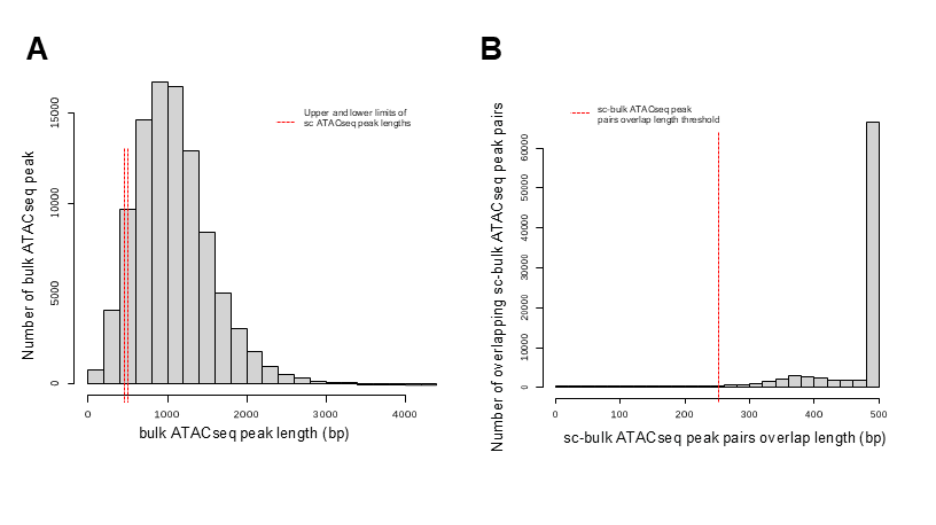

**Supplementary Figure 4. Bulk and single-cell ATACseq results in teloHAEC cells are concordant.** Open chromatin fold-changes (FC) are correlated when comparing sc (x-axis) and bulk (y-axis) RNAseq results: (**A**) teloHAEC TNFα 24 hours vs non-treated (NT) teloHAEC and (**B**) teloHAEC TNFα 24 hours vs teloHAEC TNFα 4 hours. We note that the Pearson’s correlation coefficient is markedly lower in the TNFα 24 hours vs TNFα 4 hours comparison (*r*_24h_4h_=0.13 vs *r*_4h_NT_=0.60 and *r*_24h_NT_=0.59).Given the specificities of our ATACseq experiments, the bulk ATACseq protocol detects DOP with smaller FC in teloHAEC: (**C**) TNFα 4 hours vs NT, (**D**) TNFα 24 hours vs NT and (**E**) TNFα 24 hours vs TNFα 4 hours.

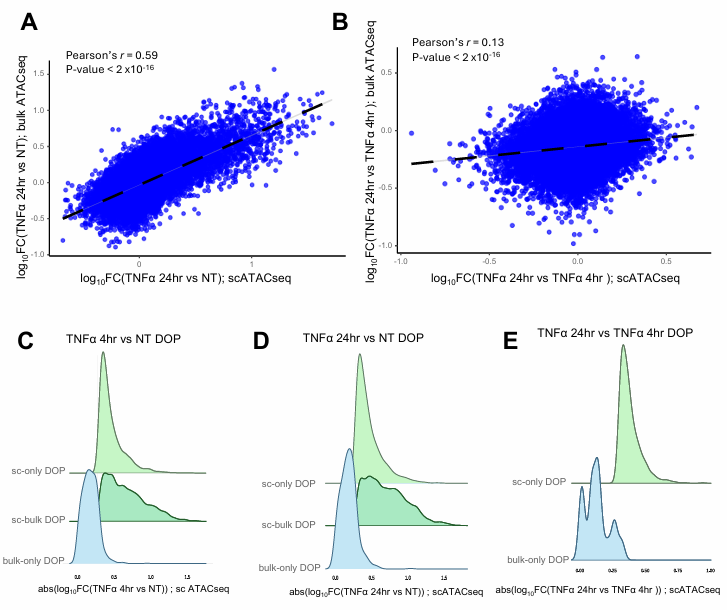

**Supplementary Figure 5. Identification of regulatory links between candidate cis-regulatory elements and genes in teloHAEC using bulk and sc data.** Bar plot summarizing the number of cCRE-gene links identified with both the bulk and sc data (ABC+scE2G), bulk only (ABC-only) and sc-only (scE2G-only) per treatment conditions. NT, non-treated.

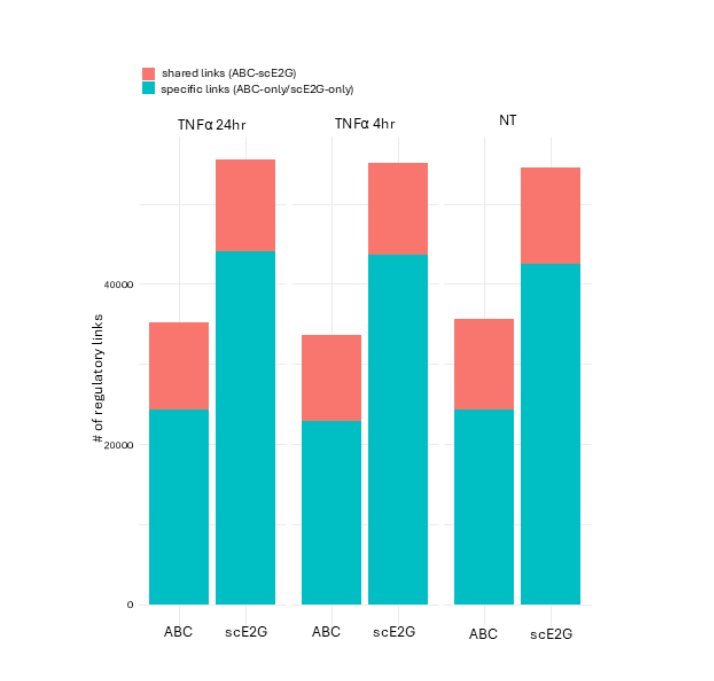

**Supplementary Figure 6. Coronary artery disease (CAD) and diastolic blood pressure (DBP) fine-mapped variants captured by teloHAEC regulatory links.**  (**A**) Jackknife estimates of the per-SNP heritability in ABC-only cCRE. The 110^th^ SNP block (chr9:20616618-33217212) is responsible for the non-significant enrichment. **(B)** Scatter plots of CAD linkage disequilibrium (LD) score regression-based heritability estimates (x-axis) for variants within five regulatory link categories, along with their corresponding enrichment P-values (y-axis) when locus chr9:20616618-33217212 is removed from the analysis. The ABC-only CAD enrichment becomes significant. Scatter plots of the number of **(C)** CAD and **(D)** DBP fine-mapped variants captured against cCRE genomic coverage. We define genomic coverage as the number of base pairs covered by cCRE over the number of base pairs in the human genome.

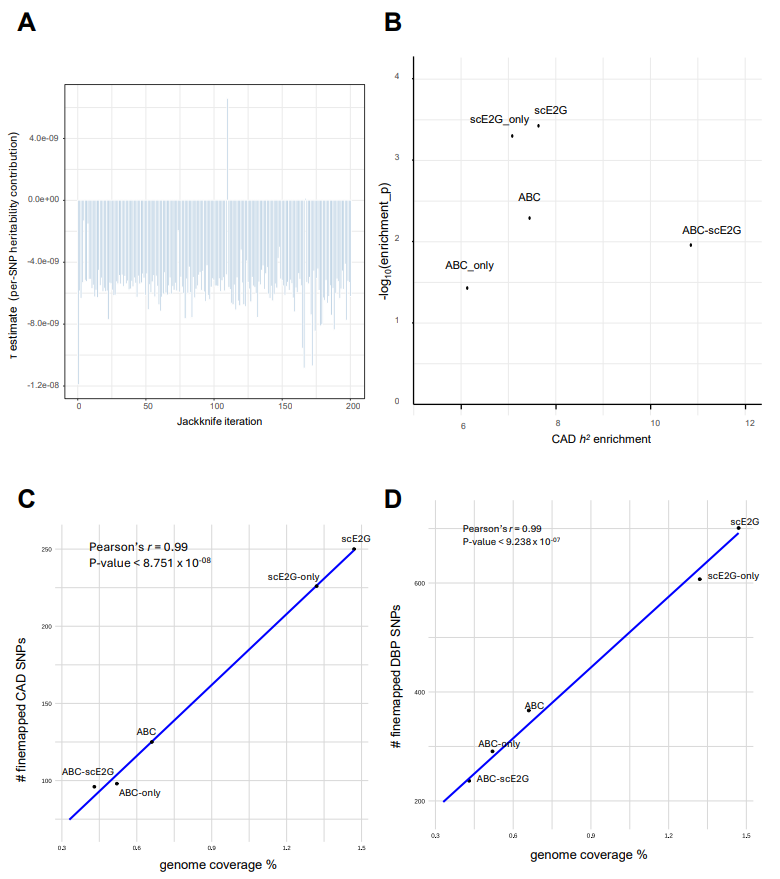
